# A mismatch between slow protein synthesis and fast environmental fluctuations determines tradeoffs in bacterial proteome allocation strategies

**DOI:** 10.1101/2025.07.22.666192

**Authors:** Leron Perez, Jonas Cremer

## Abstract

Microbes live in environments that fluctuate faster than they can adjust their cellular machinery. To survive these fluctuations, they must dynamically regulate protein synthesis—a resource-intensive process that is often slower than environmental changes. Here, we develop a mechanistic model coupling antibiotic kinetics with dynamic proteome allocation to understand how limitations in translational capacity shape acclimation strategies. Using translation-inhibiting antibiotics and resistance proteins, we show that the temporal mismatch between environmental perturbations (seconds) and protein synthesis responses (hours) creates a growth advantage for anticipatory strategies where cells pre-synthesize resistance proteins before antibiotic exposure. Further, we find that the largest benefits of anticipation and the largest protein fractions reserved for anticipation are realized in environments with multiple antibiotics, suggesting that anticipation is most important in complex environments. This work establishes a framework for quantifying the costs and benefits of various acclimation strategies in dynamical environments based on the fundamental constraints of protein synthesis, with implications for microbial ecology, antibiotic resistance, and biotechnology applications.

## I. INTRO

Microbes live in environments that change rapidly and unpredictably. To maintain energy homeostasis and biomass synthesis in these fluctuating environments, microbes dynamically adjust their cellular machinery[1– 6]. This adjustment, which we call acclimation to distinguish it from evolutionary adaptation, requires synthesizing new proteins. Acclimation is necessarily proteomic because proteins are the enzymatic engines driving the bulk of cellular processes, ranging from carbon catabolism to motility [7, 8]. But making new proteins is resource intensive [9–12] and slower than many environmental fluctuations[13, 14].

Accordingly, protein synthesis at the right time and in the right quantity is crucial [15, 16]. Too many, too few, or the wrong kind of proteins can doom a cell [17–21]. Due to the differing timescales of environmental fluctuations and protein synthesis, cells synthesizing proteins based solely on their current environment may perish. For example, a bacterium suddenly exposed to toxins often will not have enough time to synthesize resistance proteins before the toxin does its harm, making preemptive synthesis of resistance proteins essential. On the other hand, producing such proteins continuously when they are not needed is costly, as it diverts limited cellular resources [22].

Previous studies on acclimation strategies have considered specific cellular phenotypes which promote or hinder bacterial growth and survival depending on the encountered environmental conditions [23, 24]. The simplest strategies are constitutive or responsive, meaning proteins are either synthesized at a constant rate or synthesis is induced by environmental cues [25–27]. Bet hedging strategies rely on stochastic gene expression to generate a heterogeneous population [28–30]. This heterogeneity can be advantageous in fluctuating conditions because some cells are equipped to cope with future fluctuations [1, 31]. More recently, anticipatory strategies, in which microbes pre-synthesize proteins in expectation that the environment will change soon have been studied [2, 3, 32]. Studies on anticipation have particularly highlighted the tradeoff between maximizing current growth and adaptability to future conditions in fluctuating environments [33–37].

Since the speed of microbial acclimation is tied to protein synthesis rate, which in turn depends on the current proteome and the cell’s environment, further elucidating how microbes acclimate to fluctuating conditions requires studying the feedback between growth and the dynamic regulation of protein composition that shapes it.

To capture the physiological details that define the major constraints and tradeoffs inherent in acclimation, a useful theory of growth must, in our opinion, describe how protein synthesis and proteome composition translate into growth without getting lost in molecular details. Here we build on proteome allocation models [7, 38–44] to study acclimation strategies. These models equate growth rate with the rate of protein synthesis which results from the fluxes through cellular metabolism and translation. By representing the proteome as a small number of protein sectors, they succeed in rationalizing cellular ribosome content and the costs of protein synthesis. However, they are limited in their capacity to explain how cells balance their myriad processes and non-ribosomal protein allocation [45]. Across microbes, proteomes consist of up to 20-30% unused proteins, something which is not satisfactorily explained by existing growth models [17, 39, 40, 46–50].

In this study, we focus on ribosome-inhibiting antibiotics and the biosynthesis of resistance proteins to investigate the role of conditionally unused proteins and the tradeoffs between different acclimation strategies. The mechanistically well-established coupling between antibiotics and protein synthesis provides an ideal test case to illustrate and explore how physiological constraints shape acclimation strategies. Building on previous phenomenological models and observations in steady conditions [51– 54], we couple a model of antibiotic kinetics with a dynamical proteome allocation model. We systematically explore how different timescales of exposure, susceptibility, and response affect acclimation strategy success and the amount of conditionally unused proteins.

## II. RESULTS

### A. Ribosome allocation describes microbial growth with antibiotics

To study acclimation strategies, we build on recent progress in dynamical protein allocation models [53, 55– 62], explicitly going beyond steady state growth and accounting for dynamical changes in protein synthesis, proteome composition, and environmental conditions. We consider a coarse-grained description of the cellular proteome divided into four sectors by mass fraction: translation, *ϕ*_*R*_; metabolism, *ϕ*_*M*_; antibiotic resistance, *ϕ*_*AR*_; and other cellular processes, *ϕ*_0_.

Growth rate, *λ*, is given by the synthesis of novel protein mass *M*. The synthesis of new protein mass depends on the mass fraction of translating ribosomes 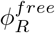 and the translation rate *γ* with which ribosomes make new proteins (Fig. 1, Eqn 1). Following the protein allocation logic, we include charged tRNA species and other needed metabolites as one effective pool of precursors, *c*_*pc*_. Precursor concentration is given by the balance of metabolic synthesis and translational consumption (Eqn 2).

**FIG. 1.**
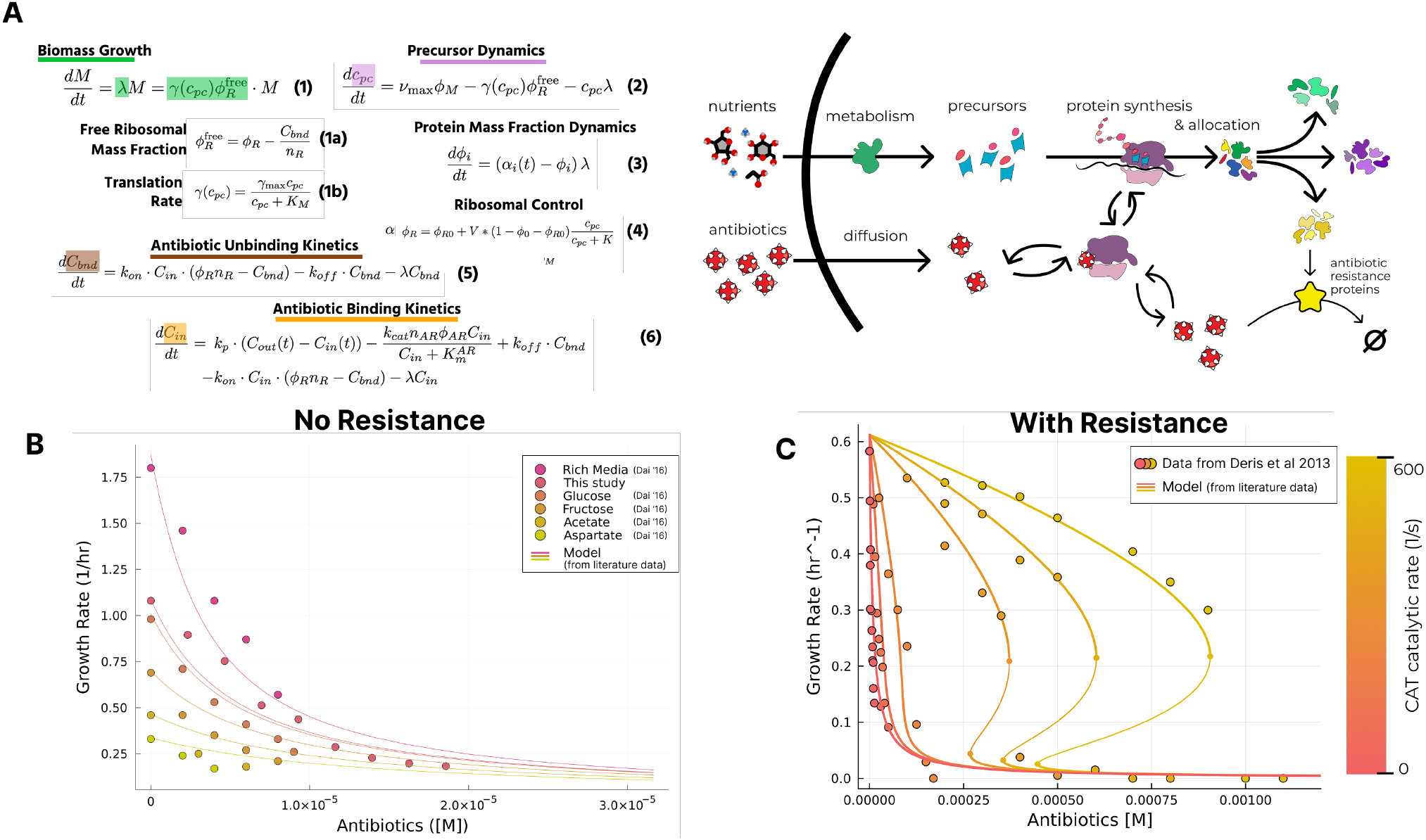
Model summary and data validation. (A) The full set of equations describing proteome allocation and growth dynamics in antibiotics and a cartoon illustrating how the modeled quantities interact. Equation 4 is parametrized to fit precursor concentrations to observed ribosome fractions during steady state exponential growth (see SI Section **??**) (B) Comparison of modeled and measured steady state growth rates in increasing antibiotics with no antibiotic resistance. Data from Dai et al 2016 [59]. (C) Comparison of modeled and measured steady state growth rates in increasing antibiotics with antibiotic resistance. Data from Deris et al 2013[51]. The model parameters are derived from literature data as discussed in the text. The metabolic rate was fit to growth with no antibiotics. With that sole exception, the model is not fit to the data.

Growth rate also sets the total amount of protein that can be divvied up by the allocation fractions, *α*_*i*_. Each *α*_*i*_ is the instantaneous fraction of new proteins allocated to protein sector *ϕ*_*i*_ (Eqn 3). In this framework, an acclimation strategy is defined by how the allocation fractions, *{α*_*i*_*}*, change over time and in response to fluctuations. For the ribosomal allocation fraction, *α*_*R*_, we employ an empirical relation (Eqn 4) motivated by past studies establishing that nutrient availability regulates ribosomal mass fraction via ppGpp sensing of uncharged tRNA levels [20, 63, 64], (see Fig. S1).

To account for translation-inhibiting antibiotics and antibiotic resistance, we explicitly model the internal antibiotic concentration as a balance of permeation into the cell, binding and unbinding to ribosomes, and degradation by the antibiotic resistance proteins, *ϕ*_*AR*_ (Eqn 5 and 6). Bound ribosomes are no longer translating, reducing the pool of active ribosomes (Eqn 1a) [53, 59]. To parametrize the model, we used data from prior studies on growth and ribosome content, the antibiotic permeability across *E. coli* ‘s outer membrane, the binding kinetics of chloramphenicol, and the catalytic activity of chloramphenicol acetyl-transferase [22, 43, 55, 59, 65–79]. The model parameters and their estimation are discussed further in the supplementary.

To validate the model we first analyzed how steady growth rates vary with chloramphenicol concentrations and carbon sources in the absence of any antibiotic resistance, comparing model predictions with data from Dai et al. (Fig. 1B). In strong agreement between model prediction and data, growth rates fall rapidly as the chloramphenicol concentration increases. We additionally validated our model in a dynamic chloramphenicol shock. *E. coli* growing at steady state in glucose minimal media were subjected to a 5 *µ*g/mL challenge. Our data shows that *E. coli* dynamically increases ribosomal mass fraction in response to an antibiotic shock with the model capturing the trend (see Fig S3).

Next, we compared our model to the steady-state growth rates of resistant microbes constitutively expressing chloramphenicol acetyl-transferase (CAT) [51] (Fig 1C). With the model parameters derived from literature, our mechanistic antibiotic dynamics strongly agrees with the data and captures the same bistability modeled phenomenologically in Deris et al [51]. Our only assumption was that *ϕ*_*AR*_ = 0.01 which was the maximal value of antibiotic expression for which our model matched the normalized growth in the absence of antibiotics presented in Deris et al. In our model, the collapse in growth arises from an overabundance of precursors leading to increasing fractions of the proteome dedicated to ribosomes at the cost of all other protein fractions. At a critical threshold, the diminished levels of constitutively expressed resistance are no longer sufficient and the fraction of total ribosomes spikes to compensate for all of the ineffective bound ribosomes. This results in a sharp drop in growth when sufficient levels of resistance are expressed (Figure 1C).

In sum, the comparison of model predictions and experimental observations in steady conditions confirms the model’s parametrization and its ability to describe major growth phenotypes.

### B. Anticipation balances protein maintenance cost by enabling a fast acclimation to antibiotics

We next consider a sudden up-shift in antibiotics in which exponentially growing cells are subject to a subinhibitory antibiotic shock of 5 *µ*g/mL (*≈* 15*µM*) chloramphenicol. To assess the growth effects of different acclimation strategies, we modeled a susceptible strategy with no resistance (*α*_*AR*_ = 0) and strategies with different antibiotic resistance allocations.

In the susceptible strategy, the intracellular antibiotic concentration spikes seconds after the shift, faster than the cell can mount any coordinated protein response. Ribosome binding kinetics happen at a similarly fast time scale, so the growth rate plummets as well during this immediate shock (Fig 2A, 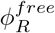 line and growth rate graph). After that, the cell begins adjusting. Precursors rise, indicating there is not enough biosynthetic capacity (Fig S4, upper row). More ribosomes are made, albeit at a reduced effectiveness since many of them are immediately bound by chloramphenicol (Fig 2A, 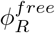 and *ϕ*_*R*_ lines). The new ribosomal proteins are synthesized instead of metabolic proteins, which results in a two-pronged reduction of the precursor concentration; both the supply of and demand for precursors is reduced (Fig S4, upper row). Eventually, the cell reaches a new equilibrium where it is making an increased ribosomal fraction to compensate for the antibiotic load.

**FIG. 2.**
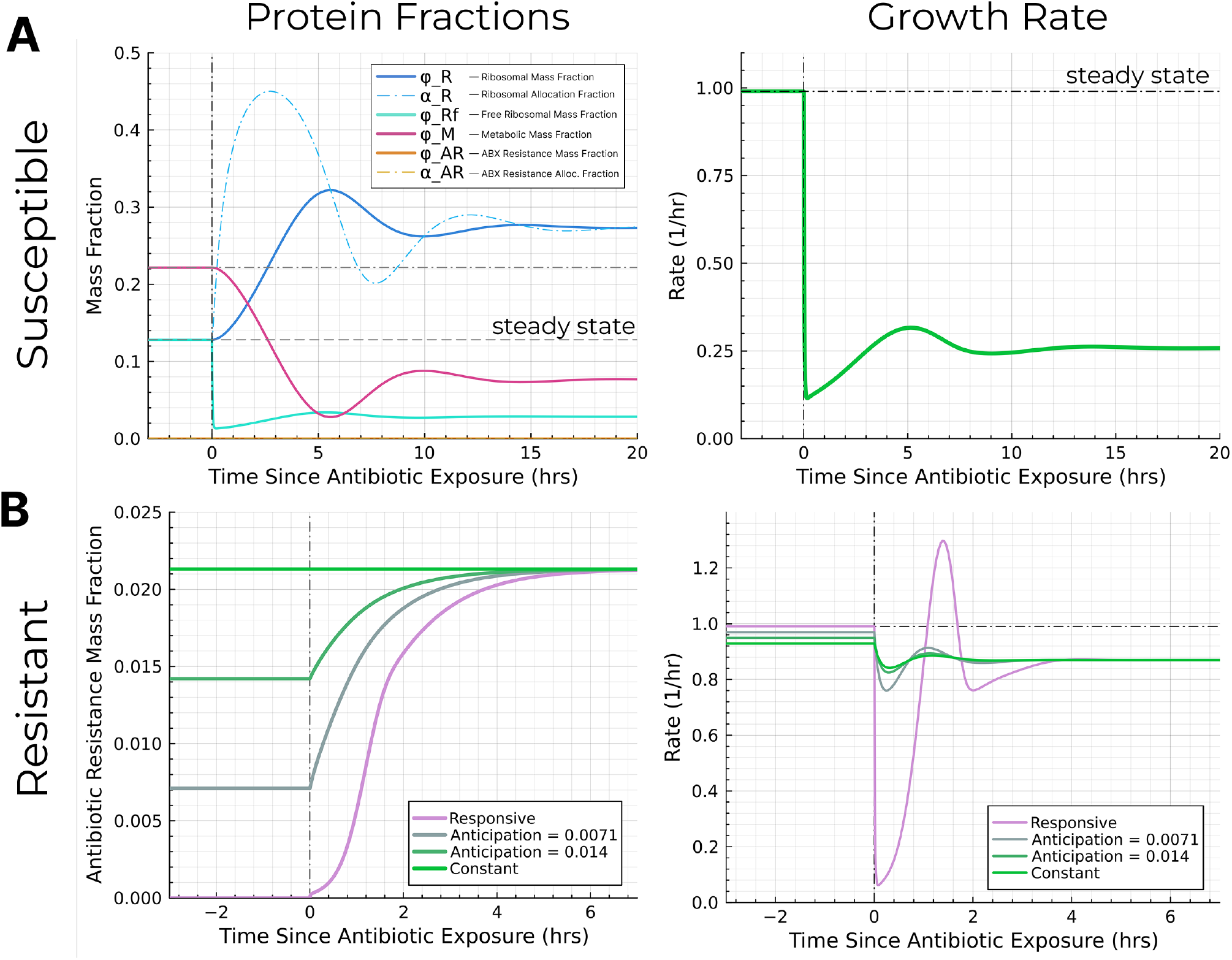
Dynamic simulations of the model in response to a step change in antibiotics. (A) The response of the susceptible strategy to a sudden 5 *µ*g/mL chloramphenicol shock, showing proteome mass-fractions, instantaneous allocations, and growth rate over time, (B) The anticipatory mass fraction and growth dynamics for a range of different resistance acclimation strategies

Next, we consider how antibiotic resistance can boost growth during the shift. We examined three kinds of resistance allocation strategies, each characterized by two numbers: the allocation to antibiotic resistance in the presence and absence of antibiotics, 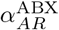 and 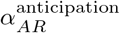. For the given external antibiotic concentration, we chose 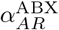 to optimized steady growth in the presence of antibiotics. The strategies differ only in 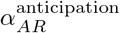. A responsive strategy does not express any resistance proteins until after antibiotics are introduced, 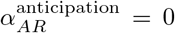. A constant strategy always expresses the same amount of resistance, 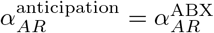. The anticipatory strategies span the space between the other two strategies, where 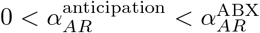.

For the responsive strategy, there is also an initial drop in growth like in the susceptible case. However, over a course of a few hours, the synthesis of resistance proteins kicks in and a substantially higher steady-state growth is achieved (Fig 2B, responsive line). In the case of a constantly expressed antibiotic resistance, the initial drop in growth is substantially reduced (Fig 2B, constant line). These benefits come at the cost of a lower steady state growth rate when there are no antibiotics. However, the decrease is relatively small and a comparatively large benefit is reaped from antibiotic resistance for a small investment cost – in steady state, antibiotic resistance maintained at a growth defect of −0.01/hr yields a growth advantage of +0.64/hr in antibiotics. The constant strategy also re-achieves its steady-state growth rate faster than the responsive strategy.

The anticipatory approach enables a smooth traversal of the constant and responsive strategies. A small additional amount of pre-synthesized antibiotic resistance significantly reduces the fitness defect from a sudden antibiotic shift (Fig 2B, anticipatory lines). This small pre-expression significantly stabilizes translation and the cell is able to cope with a sudden shock of antibiotics almost as well as a fully prepared constant resistance expression strain while also minimizing its maintenance cost.

### C. The comparative growth advantage of anticipation

To further explore the costs and benefits of anticipatory protein synthesis across different environmental conditions, we simulated growth dynamics in periodic antibiotic shocks of 15 *µ*g/mL chloramphenicol (*≈* 46*µM*). Each fluctuating condition is characterized by a period and a duty fraction, i.e. the fraction of time antibiotics are present during each cycle (Figure 3A).

**FIG. 3.**
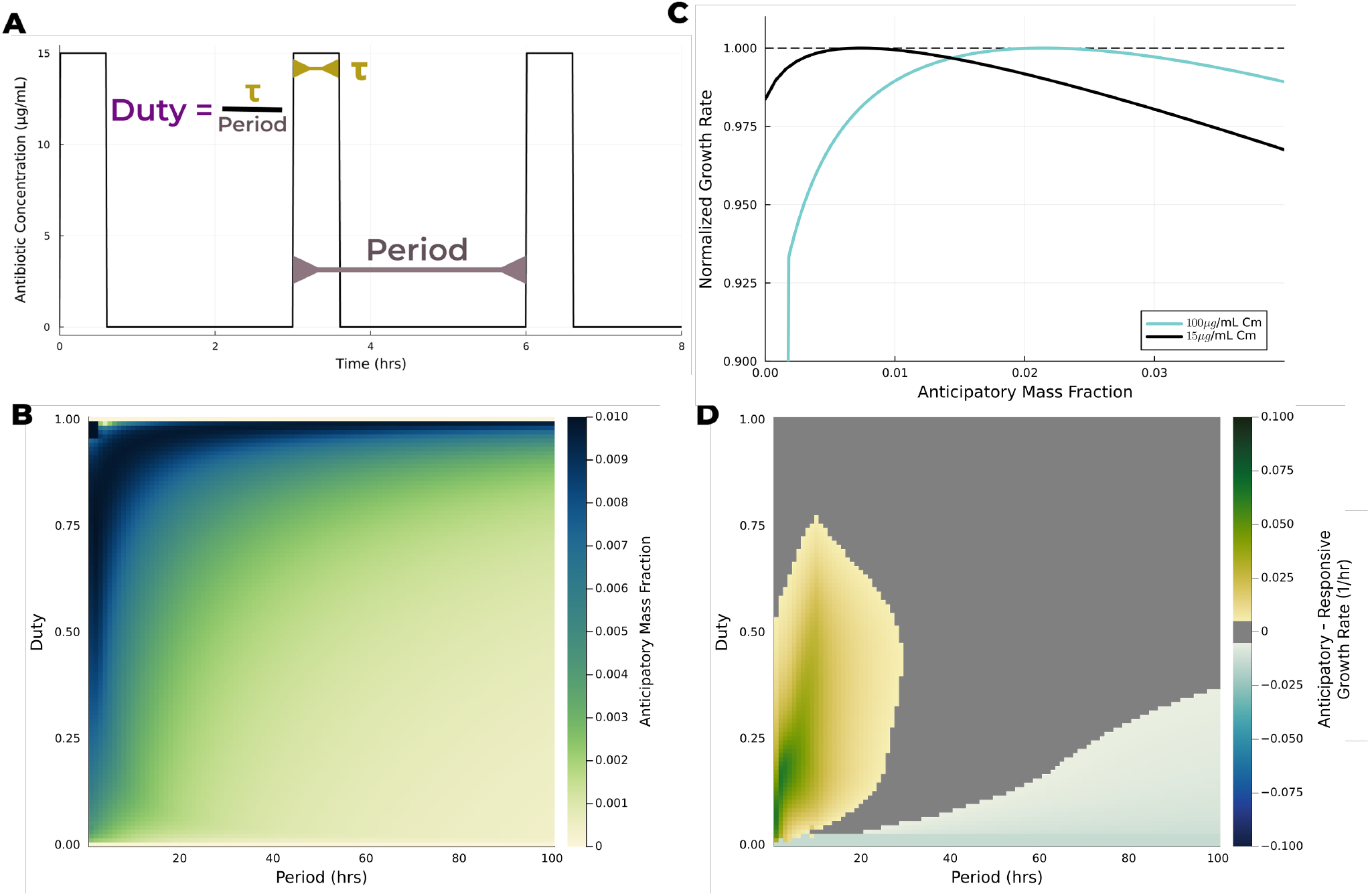
Phase diagrams illustrating the comparative benefits of strategies across various fluctuations. (A) Each antibiotic fluctuation is parametrized by a duty, a period, and by the maximal antibiotic concentration. (B) Phase diagram of the optimal protein fraction maintained for antibiotic resistance in varying fluctuations. (C) Growth rate versus anticipatory pre-expression for a given fluctuation (period = 5 hours, duty = 0.5). There is a clear optimal 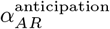 and it increases with antibiotic concentration. (D) Phase diagram of the growth difference between the anticipatory and responsive strategies (*λ*^anticipatory^ *− λ*^responsive^). A positive value means anticipation is better. For approximately neutral growth differences (*<* 0.005*/*hr) we plot a null gray value. The antibiotic concentration used was 15*µ*g/mL for all phase diagrams.

To identify the optimal acclimation strategy for fluctuating antibiotics, we simulated the growth of strategies with differing amounts of anticipatory pre-expression,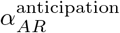, across different fluctuations. The growth rate for each condition was measured by taking the average of growth rate over a single cycle once the cell state had equilibrated to the fluctuating condition. We then computed the optimal anticipatory mass fraction for each fluctuating condition and found that optimal anticipation varies strongly with the period and duty of fluctuations (Figure 3B). Notably, the responsive strategy 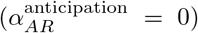 is only optimal at the duty=0 boundary (no antibiotics). Everywhere else, some minimal amount of pre-expression provides a fitness benefit. At the duty = 1 boundary, anticipation has no impact on growth rate, since the microbes are always exposed to antibiotics. At short periods and high duty, anticipation also does not affect growth much because the protein composition does not have enough time to change significantly.

However, while 15*µ*g/mL is bactericidal for nonresistant strains [80], microbes exposed to antibiotic fluctuations are likely to experience higher doses, even beyond 250*µ*g/mL [81]. Higher doses results in higher fractions of the proteome devoted to anticipation since growth peaks for a higher value of anticipatory mass fraction in Figure 3C as the antibiotic concentration rises. Another interesting aspect of the relation between growth rate and anticipation is that over-anticipating is much less costly than under-anticipating. Asymmetric selective pressures will lead anticipation to increase to an optimum faster than it decreases to an optimum (Figure 3C). In fluctuations with variable levels of antibiotics, we thus expect the optimal anticipation level to approach the optimal anticipation for the highest level of antibiotics encountered, rather than the average of encountered concentrations.

Until now, we considered optimal anticipation in different antibiotic fluctuations. Yet for a given microbe, anticipation is fixed and takes evolution to change. Therefore, we analyzed the growth advantage a given anticipatory strategy 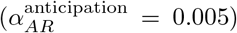 has over the responsive strategy 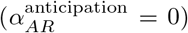; first by simulating their average growth rates over each fluctuation condition (Fig. S5) and then by subtracting the responsive growth rates from the anticipatory ones (Figure 3D).

At duty=0, there are never any antibiotics and correspondingly, this is the largest negative difference, because the maintenance cost of anticipation never pays off. At the opposite duty=1 boundary, there is a constant antibiotic exposure and the two strategies are identical, by construction. As the period of fluctuations gets long, the fitness contribution from steady-state growth rates dominates, and so for periods *>* 20 hrs, the growth rate converges to a duty-weighted average of the two steady-state growth rates (Fig. S5).

The benefit of anticipation comes at shorter times. There is a large, fixed fitness cost that the responsive strategy bears every time the environment transitions to antibiotics. For very short fluctuations (approximately, period *∈*(0, 5] hrs), the responsive strategy is less deleterious (Fig. S5). For an intermediate timescale (approximately, period *∈* [5, 15] and depending on the duty), the responsive strategy bears the full fitness cost of the antibiotic exposure every cycle, and anticipation has the biggest advantage. As the period gets longer, the maintenance cost of the pre-expression slowly catches up to the benefit of a faster transition. Notably, we likely overestimate the maintenance cost that cells experience, meaning that real microbes probably anticipate more and for longer. Conceptually, maintenance costs arise from making useless proteins at some fixed rate. Maintaining the same mass fraction of useless protein at a faster growth rate necessitates a higher rate of useless protein synthesis than at slower growth rates. Consequently, in real settings, where growth is not maintained at the consistently high levels that we modeled, maintenance costs are lower and we expect more anticipation to persist.

In summary, anticipatory acclimation leads to faster growth than responsive acclimation across a broad range of antibiotic fluctuation conditions. Anticipation is most advantageous for intermediate fluctuation timescales where the fitness benefit of rapid acclimation outweighs the costs of pre-expression maintenance due to the high frequency of transitions.

### D. Anticipation facilitates acclimation to the compounded stress of multiple simultaneous antibiotics

We expect anticipation to be especially relevant in complex natural environments where microbes are exposed to multiple, fluctuating stressors. To illustrate this we consider multiple translation-inhibiting antibiotics and explore how different acclimation strategies fare against simultaneous or overlapping antibiotic shocks. We extended our model by considering *n* antibiotic resistance fractions,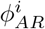, that each degrade a different antibiotic. Each antibiotic targets an independent site on the ribosome in our model. The ribosome has at least 13 independent antibiotic binding sites [82], so there is ample potential for a high anticipatory load on account of different antibiotics. Furthermore, each resistance pathway has its own regulatory strategy; bacteria might employ an anticipatory strategy for one antibiotic and a responsive strategy for another.

To explore the costs and benefits, we assumed the kinetics of antibiotic inhibition and resistance are identical for all antibiotics and their resistance protein fractions. Additionally, we considered a fixed set of fluctuations with a period of 3 hours and a duty of 0.2. Figure 4A shows an example of growth dynamics for a strain exposed to two antibiotics with no resistance. We first compared the growth advantage of two anticipatory strains; one exposed to a single antibiotic at 15*µ*g/mL, (Fig 4B, dashed line) and another exposed to 4 antibiotics, each at 3.75 *µ*g/mL for an equivalent total of 15*µ*g/mL (Fig 4B, black line). We observe that the benefits of anticipation are more significant and peak at a higher level of anticipation in response to multiple antibiotics.

**FIG. 4.**
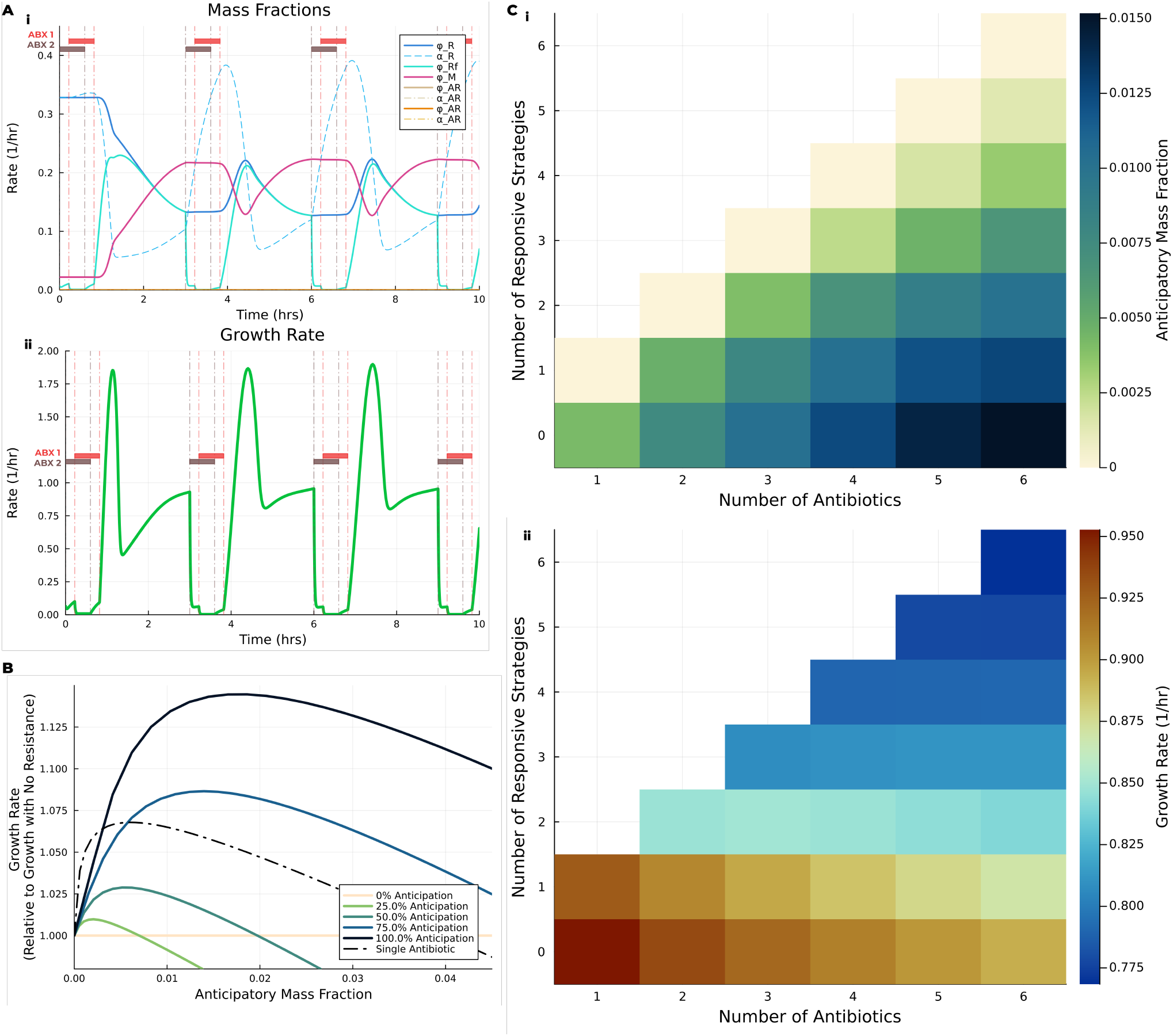
Acclimation in response to multiple antibiotics. (A) Dynamical simulations of our model extended to two antibiotics in fluctuations with a period of 3 hours and a duty fraction of 0.2. Each antibiotic binds the ribosome at independent sites and no antibiotic resistance is expressed. (B) A strain with four resistance protein fractions where each fraction is either anticipating or responding to four simultaneously applied antibiotics. A dashed line showing the growth behavior of a strain exposed to a single antibiotic and expressing the corresponding resistance is shown for comparison. (C) Increasing antibiotics leads to an increasing fraction of the proteome devoted to anticipation, (i) illustrates the optimal total anticipatory mass fraction and (ii) illustrates the growth rate attained at the given anticipation.

We then compared the growth effects of different combinations of regulatory strategies for bacteria exposed to simultaneous antibiotic shocks. We considered 0-4 anticipatory strategies in bacteria with 4 resistance protein fractions. Any non-anticipatory resistance was responsively regulated. The anticipatory mass fraction expressed by each strain was equally spread among the anticipating resistance fractions. The growth benefic increased with anticipation, with the largest benefit coming from 100% anticipation (Fig 4B) which also employs the highest levels of pre-expressed resistance. In nonsimultaneous exposures, the exact benefit of each anticipatory fraction further depends on the degree of overlap or separation of each shock and further the order of the shocks matters.

More anticipation is needed as the number of antibiotics increases. To show this, we extended the simulations from Figure 4B to find the optimal anticipation and the resulting growth rate for varying combinations of strategies and the number of simultaneously-occurring antibiotics (Figure 4C). As the number of antibiotics increases, growth rate falls even though the total antibiotic concentration is held constant at 15*µ*g/mL (panel C ii). For a given number of antibiotics, growth is always maximized by pre-expressing all of the resistance proteins (e.g. growth is maximal in the bottom row where all strategies are anticipatory). Furthermore, the optimal anticipation increases with the number of simultaneous antibiotics.

## III. DISCUSSION

Microbial survival in changing environments depends on an appropriate regulation of protein synthesis [6, 20, 34, 57, 58, 83, 84]. Understanding cellular physiology beyond tightly controlled steady lab environments, whether in their native niches or in industrial fermenters, requires models that closely couple acclimation strategies with detailed considerations of protein synthesis and microbial growth [85]. By using the mechanistically well-established connection between translation-inhibiting antibiotics and protein synthesis [54, 82], we demonstrated how different protein synthesis strategies determine growth across a variety of fluctuating environments. We specifically focused on anticipatory pre-expression of antibiotic resistance proteins and showed that anticipation increases microbial fitness across a broad range of fluctuations.

The physiological basis for the growth advantage of pre-expression lies in the temporal mismatch between environmental perturbations (fast) and the cellular responses involving protein synthesis (slow) [86, 87]. Chloramphenicol permeates the cell and binds ribosomes within seconds, while protein synthesis can require hours to significantly alter proteome composition [72]. While we focused on translation-inhibiting antibiotics and specifically on chloramphenicol, this asynchronicity is widespread across the microbial world, from soildwelling photoautotrophs [88] to marine heterotrophs and beyond [90]. Different kinetics of stress and response lead to different optimal strategies [32, 91], but the mismatch between the timescales of environmental change and microbial response is the rule, rather than the exception. Accordingly, microbes across all sorts of environments must find ways to bend their too-slow protein synthesis to surf the waves of change [92, 93].

This need for matching protein synthesis to fluctuations is especially vital in complex environments, where multiple simultaneous stressors can compound the need for efficient acclimation [94–96]. For example, our results show that anticipation is even more important when microbes are exposed to multiple antibiotics, as opposed to a single antibiotic. Furthermore, the model predicts that microbes should err on the side of over-anticipation in less predictable environments, since insufficient anticipation has a much larger growth disadvantage than over-anticipation.

While anticipation is beneficial, it necessarily requires the maintenance of temporarily unutilized proteins. Our work taken together with the observation that microbes maintain up to *∼*30% of their proteomes as apparently unutilized proteins[46, 47, 49, 50, 87, 97] suggests that anticipating complex, fluctuating environments strongly shapes the physiology and evolution of cells. Further study of these unutilized proteins could better inform our knowledge of natural environments and microbial acclimation to the fluctuations within them.

Further research can build on our protein allocation framework with anticipation to quantitatively infer what environments a microbe likely inhabits. However, while microbial proteomes might capture certain characteristics of complex fluctuating environments, they do not provide a perfect record. Particularly, fluctuations of universal stressors like certain antibiotics or temperature might leave widespread footprints in proteomes. But environments like the gut or tropical soils contain a myriad of microbes that specialize to different carbon sources each with their own unique fluctuations. As a result, we expect microbes in the same environment to contain distinct anticipatory signatures.

Notably, the study of acclimation via protein allocation models also implies that cells’ finite protein capacity promotes diversification [37]. It has not been clear why a strain can not just switch between an infinite number of environment-specific optimal pathways. By motivating that strains are limited to specializing in a limited number of environments due to the proteomic load of anticipatory proteins, dynamical protein allocation models like ours provide a framework to quantitatively understand growth dynamics in diverse microbial communities. Future research can explore how strains within diverse communities allocate their proteome in fluctuating environments. Future work could also explore the costs of regulation and lightly utilized genes to understand how specialization emerges on longer timescales, perhaps by quantifying how long an unused responsive strategy might persist in a microbial genome.

Finally, biotechnology could also benefit from further efforts to quantify the coordination of protein synthesis in changing environments. For industrial fermenters hundreds of thousands of liters in volume, heat and mass transport cannot keep up with microbial growth even with the help of active stirring. As a result, microbes experience large temperature and chemical fluctuations[94, 98]. Better understanding how protein synthesis is coordinated to changing conditions could inform more robust production strains and perhaps even bioreactors codesigned with the microbes they are meant to grow.

## ACKNOWLEDGMENTS

We thank members of the Cremer lab for valuable discussions and comments on the manuscript, Eva Brand for comments and feedback on the manuscript, and Drew Endy for suggesting we focus on antibiotics and earier commentary. This work was supported in part by the NIH Molecular Biophysics Training Program (T32 GM136568) and a Terman Fellowship from Stanford University.

## DATA AVAILABILITY

Code and data are available at this link.

