## Supplemental Info for "A mismatch between slow protein synthesis and fast environmental fluctuations determines tradeoffs in bacterial proteome allocation strategies"

(Dated: July 22, 2025)

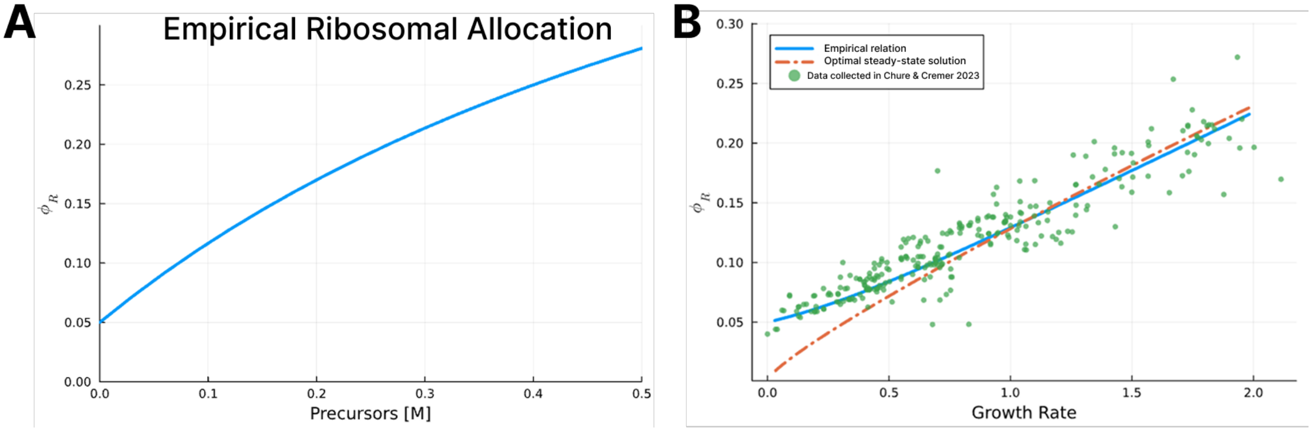


FIG. S1. To model the allocation of protein mass to ribosomes we modeled the allocation to ribosomal mass fraction, $\alpha_R$, as a function of the precursors, $c_{pc}$, in line with the dependency of ribosomes on ppGpp and the ratio of charged to uncharged tRNA [1–3]. This figure shows how, at steady-state ($\alpha_R = \phi_R$), our empirical relation (equation S4) captures the observed growth behavior. Data is from the collection of sources aggregated in [4]. Equation S4 is parametrized by the data depicted; we get $V = 2$ and $K = 0.8$ with the constraint that $\phi_{\mathrm{R0}} = 0.05$. (A) shows the relation of precursors to the ribosomal mass fraction as defined by equation S4. (B) compares the growth behavior resulting from equation S4 to data

### S1. EXPERIMENTAL METHODS

The following methods detail the experiments that were conducted to validate the model. The steady state growth data is shown in Fig 1B. The dynamical antibiotic shock growth and ribosome fraction data is shown in Fig. S2.

#### 1. *Steady State Growth*

*E. coli* K-12 substrain NCM3722 was grown in an LB seed culture overnight or until OD began saturating ($\sim 0.2 - 0.4$). For each antibiotic concentration, the culture was diluted into a pre-culture tube containing the appropriate chloramphenicol concentration in $\sim$ 3mL aerobic glucose minimal media (GMM) at 250 RPM in an orbital water bath shaker. Carbon and nitrogen in GMM was 10 mM glucose, 10mM ammonium chloride. A 500mL 4x N-C- stock was prepared with 2g $K_2SO_4$, 27g $K_2HPO_4$, 9.4g $KH_2PO_4$, 0.8 mL of 1M $MgSO_4$, and 5g $NaCl$, filled to 500 mL with double-distilled water. The pre-cultures were grown

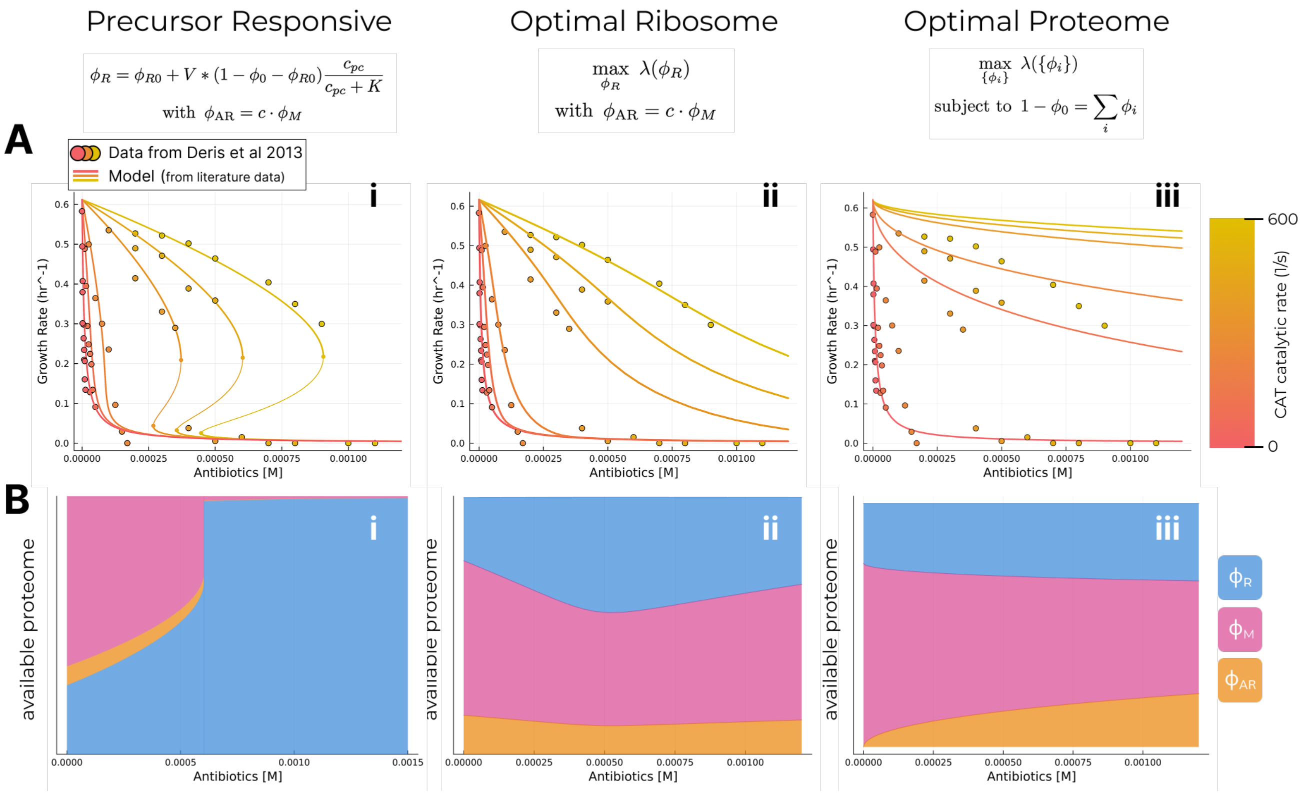


FIG. S2. In this figure, we model several different regulatory schemes for $\alpha_R$ and $\alpha_{AR}$. In the main text, the model and data match best for the "Precursor Responsive" regulation and these are the relations used in the models presented in the main text. The antibiotic resistance was expressed constitutively in Deris et al [5]. (A) depicts the relationship between growth rate and antibiotic concentration for several levels of antibiotic resistance. (B) shows cartoons depicting how the protein mass fractions shift for each regulatory scheme as antibiotics increase. The optimal proteome allocation results in a proportional increase of resistance as antibiotics increase, with a relatively small increase in $\phi_R$. We expect that antibiotic inducible promoters (i.e. pTet) can achieve this optimal proteome allocation. The optimal ribosomal allocation supposes that $\phi_{AR}$ is a fixed fraction of $\phi_M$. As a result, as antibiotics increase there is a critical point where increasing $\phi_R$ to compensate for bound ribosomes becomes detrimental because it diminishes $\phi_{AR}$ as well. Lastly, for the precursor responsive strategy, $\phi_R$ increases as the imbalance between metabolic and biosynthetic fluxes increases. In antibiotics, this leads to a runaway feedback that results in a critical point where $\phi_R$ increases to the exclusion of all else.

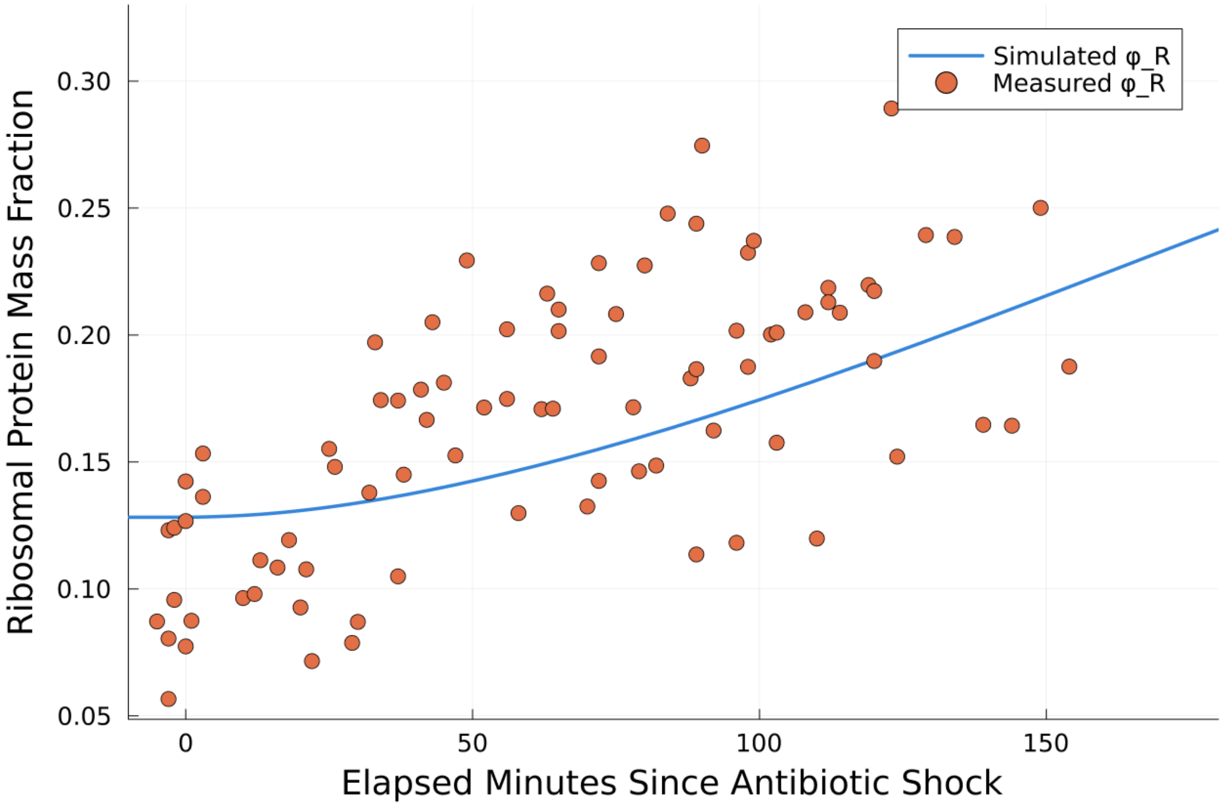


FIG. S3. A comparison of experimental $\phi_R$ dynamics with our modeled dynamics in response to an antibiotic shock. As described in sections S1 2 and S1 3, we exposed *E. coli* to an antibiotic shock and then quantified how its ribosomal mass fraction adjusted by measuring the total RNA and protein content of cell extracts spectrometrically. While there is agreement between the general trend of the model and the data, the data is too noisy for firm conclusions. We refrain from averaging or fitting any curves to the data because we believe this would be misleading and that more precise measurements of $\phi_R$ are needed to assess the agreement or disagreement of the model and data. Nonetheless, others have reported the same predictions of damped oscillations[6]; in antibiotics we expect the oscillations would be especially pronounced. Others have experimentally observed such oscillatory dynamics arising from synthetic circuits [7].

for at least 10 doublings and then re-diluted into an identical glucose minimal media with antibiotics. Pre-culture was maintained in steady-state and diluted as necessary to ensure the culture did not saturate. OD measurements were recorded for each culture. Growth rate was calculated by a linear regression to the log OD for ODs in the linear range of the spectrometer (0.04 - 0.4 OD).

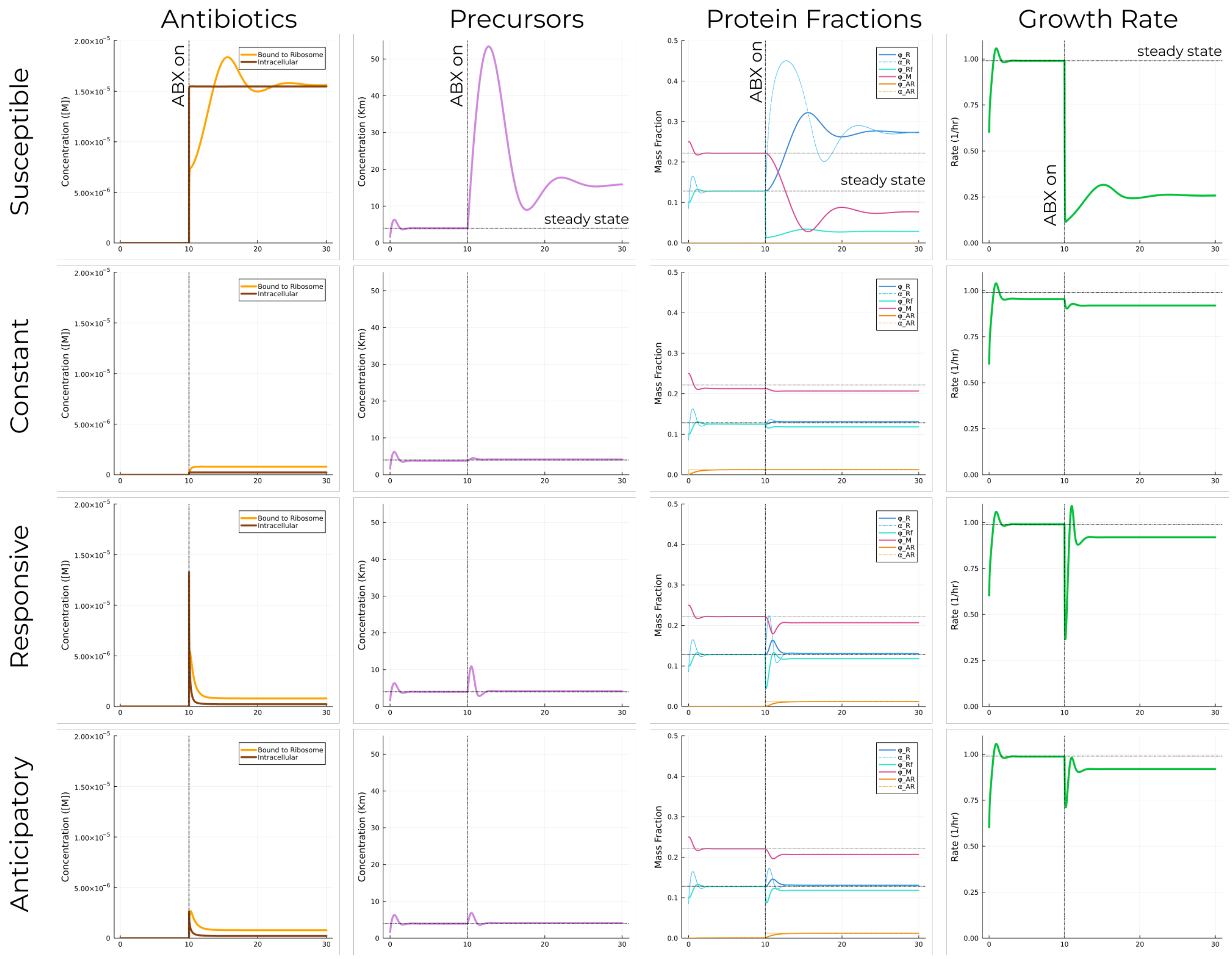


FIG. S4. A full panel of the model's output for each of the different acclimation strategies. Across all strategies, the antibiotics rapidly equilibrate and then protein synthesis begins responding. The susceptible strain has long lasting oscillations in proteome content, precursor concentration, and its growth rate. All the other strategies acclimate much faster and more stably to the antibiotic shock. The anticipatory strategy interpolates between the constant and the responsive strategies; jointly minimizing the cost of maintaining resistance and the time to acclimate to an antibiotic shock.

#### 2. *Antibiotic Shock Assay*

*E. coli* K-12 substrain NCM3722 was grown in an LB seed culture as in the steady state growth experiments. The culture was serially diluted into glucose minimal media (without any antibiotics) and grown overnight. The culture, still in exponential growth, was diluted

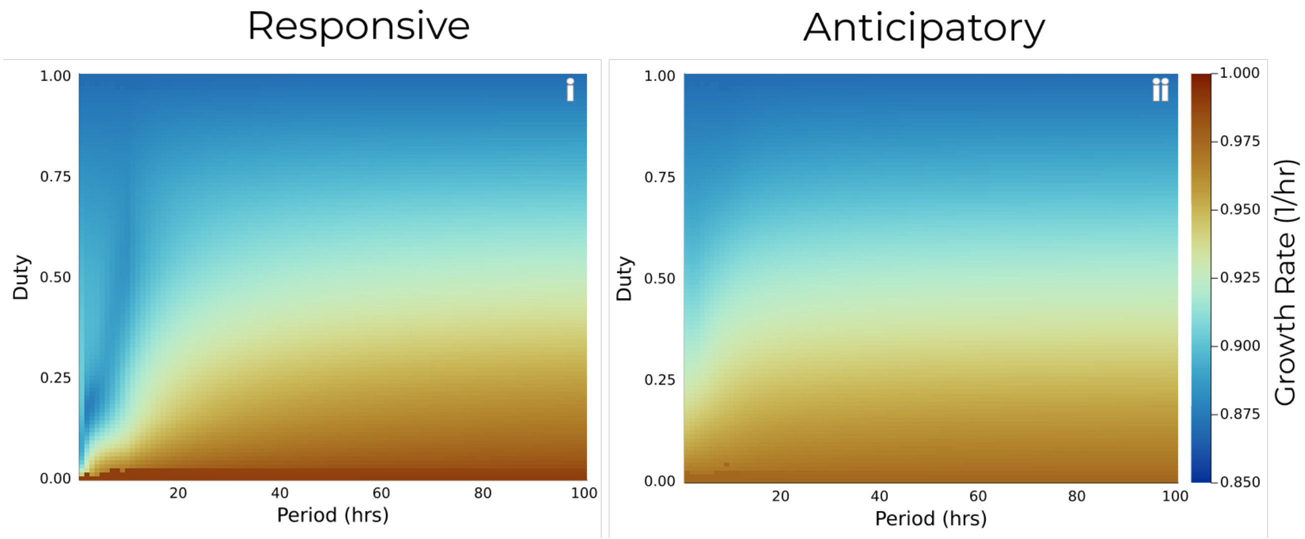


FIG. S5. Phase diagrams showing the growth rate for the anticipatory and the responsive strategies. Each pixel is the average growth rate over a full cycle once the growth rate dynamics have stabilized. For long periods the growth rates converge to the duty-weighted average of the steady-state growth rates defined by resistance expression with and without antibiotics (set by $\alpha_{min}$ and $\alpha_{max}$). At duty= 1, the growth rates are identical because $\alpha_{max}$ is the same for both strategies. They are distinguished by the differences at intermediate duties and shorter periods where the anticipatory strategy provides the largest growth benefit.

into a 140mL culture in an Erlenmeyer flask. 10 mL of 75 $\mu$g/mL chloramphenicol was dosed in to the main culture to get start the antibiotic shock, for a final concentration of 5 $\mu$g/mL. OD was measured in a quartz cuvette using 400 $\mu$L of culture. Two 1 mL biomass samples were collected for later RNA and protein extractions at each time point. One sample was collected prior to the antibiotic shock, and the remaining samples were collected in intervals of 5-10 minutes, starting immediately after the shock. Samples were immediately spun down, the supernatant was discarded and then they were put on ice. Samples were then transferred to a -80 fridge as soon as possible.

#### *3. RNA and Protein Extractions*

The RNA extraction protocol is modified from Benthin et al [8]. Cells were thawed and then washed twice in 600$\mu$L of cold 0.7M $HClO_4$ at the max centrifuge speed for 3 minutes.

The centrifuge was cooled to 4C for the duration of the extraction. After each wash, the supernatant was retained to quantify lost biomass using OD600 readings. After the second wash, cells were digested with 300$\mu$L of cold 0.3M KOH for 60 minutes at 37C. Then, we added 100$\mu$L cold 3M $HClO_4$ and centrifuged at max for 2 minutes. We then collected and saved the supernatant. The extract was washed twice more with 550$\mu$L of cold 0.5M $HClO_4$, still saving and aggregating the supernatant. RNA content was determined by measuring OD260. A negative control was added for calibrating the measurements.

Following a modification of the protocol in Herbert et al [9], for the protein extraction, we added 200$\mu$L of double-distilled water and 100$\mu$L 3M NaOH to the frozen cell pellets. Cells were then incubated at 100C for 5 minutes and then cooled to room temperature for another 5 minutes in a water bath. We then added 100$\mu$L of 1.6% w/v $CuSO_4$ and let stand at room temperature for 5 minutes. The extracts were then centrifuged at max speed for 5 minutes. Protein content was determined by measuring OD555 of the supernatants and comparing it to a standard curve created used a BSA standard.

### S2. MODELING

#### A. Derivation and Equations

We lay-out a detailed explanation of our entire model, starting with the growth rate. We equate growth rate with the rate of protein biomass synthesis. The protein synthesis is the product of ribosomal translation rate and available translating ribosomes, as in prior work [4, 6, 10–16]. To focus on the dynamics of acclimation to antibiotics and not limited resources, we model *E. coli* growing in an infinite pool of saturating media. This is equivalent to assuming we continuously transfer the bacteria to fresh media while they are growing in batch culture. We focus on the increase of protein biomass because proteins drive nearly all cellular processes, directly or indirectly. The increase in protein biomass, $M$, is given as

$$\frac{dM}{dt} = \underbrace{\lambda}_{\text{Growth Rate}} M = \overbrace{\gamma(c_{pc})}^{\text{translation rate}} \underbrace{\phi_R^{\text{free}}}_{\text{free ribosome mass fraction}} \cdot M \tag{S1}$$

where the free ribosome mass fraction, $\phi_R^{\text{free}} = \frac{M_R^{\text{free}}}{M}$, is the mass of free ribosomes divided by the total biomass. Translaton rate, $\gamma(c_{pc}) = \frac{\gamma_{\max} c_{pc}}{c_{pc}+K_M}$ follows Michaelis-Menten kinetics where we treat the ribosome-transript complex as the catalyst and the precursors, representing charged tRNA, as the substrate. The dependence of protein synthesis on the concentration of protein precursors incorporates the balance between protein synthesis and external nutrient digestion into our model.

The dynamics of the ribosomal mass fraction are a balance between newly allocated ribosomal proteins and the loss of proteins to dilution as biomass growth continues:

Ribosomal allocation fraction

$$\frac{d\phi_R}{dt} = \left( \alpha_R(t) - \phi_R \right) \lambda \tag{S2}$$

In addition to the ribosomal fraction, $\phi_R$, we model the metabolic mass fraction, $\phi_M$, the antibiotic-resistance mass fraction of proteins $\phi_{AR}$, and a static mass fraction $\phi_0$. In principle, all the protein mass fractions follow this same balance between allocation and dilution. However, we only need to additionally specify $\phi_{AR}$, since $\sum_i \phi_i = 1$ and $\phi_0$ is a constant, so with $\phi_R$ and $\phi_{AR}$ defined, we get $\phi_M$. Correspondingly, the dynamics for $\phi_{AR}$ are:

Allocation to resistance proteins

$$\frac{d\,\phi_{AR}}{dt} = \left( \alpha_{AR}(t) - \phi_{AR} \right) \lambda \tag{S3}$$

dilution

We do not explicitly consider protein degradation for any of the protein fractions, since it is negligible under most conditions for *E. coli* [17].

The allocation dynamics, $\alpha_R$ and $\alpha_{AR}$, provide extra degrees of freedom. As discussed in the main text, $\alpha_R$ is motivated by experimental studies of ppGpp and the empirical relationship observed between $\phi_R$ and growth rate. The equation for $\alpha_R$ is (see also Fig S1):

$$\alpha_R = \phi_{R0} + V * (1 - \phi_0 - \phi_{R0}) \frac{c_{pc}}{c_{pc} + K} \tag{S4}$$

We chose $V = 2$ and $K = 0.8$ to fit the data depicted in Fig S1, though there are other values of $V$ and $K$ that also fit the data well with the constraint that $\phi_{R0} = 0.05$.

$\alpha_{AR}$ varies depending on acclimation strategy pursued. For all strategies, $\alpha_{AR}$ is modeled as a fraction of $\alpha_M = 1 - \phi_0 - \alpha_R$ because this dynamic adjustment of antibiotics with $\phi_R$

most closely matches the available data (Fig S2). As suggested by the name, in the constant strategy, $\alpha_{AR}$ is a constant fraction of $\alpha_M$. In the responsive and anticipatory strategies,

$$\frac{d\alpha_{AR}}{dt} = \begin{cases} k_{ind}\left(1 - \frac{\alpha_{AR}}{\alpha_{\max}}\right) & C_{\text{out}}(t) > 0 \\ -k_{ind}\left(\alpha_{AR} - \alpha_{\min}\right) & C_{\text{out}}(t) = 0 \end{cases} \tag{S5}$$

The steady state value of $\alpha_{AR}$ is set by $\alpha_{\max}$ in antibiotics and by $\alpha_{\min}$ when there are none. For the responsive strategy, $\alpha_{\min} = 0$ and for the anticipatory strategies, $0 < \alpha_{\min} < \alpha_{\max}$. The speed of changing to a new protein allocation and inducing antibiotic resistance, $k_{ind}$, is of little consequence, since it is essentially instantaneous with respect to the speed of protein synthesis. Since, chloramphenicol-inducible genes are strongly induced at concentrations of 0.6 - 6 $\mu$M (0.2 - 2 $\mu$g/mL) [18, 19] and chloramphenicol rapidly permeates the cell membrane, $k_{ind} = 1000/hr$ is motivated by the speeds at which chloramphenicol reaches strongly inducing concentrations intracellularly (seconds).

Protein precursors represents the availability of charged tRNA and the concentration results from balance of precursor synthesis and protein synthesis rates. We model precursor synthesis as a Monod function, $\nu(c_{\text{ext}}) = \frac{\nu_{\max} c_{\text{ext}}}{c_{\text{ext}} + K_M^{\text{ext}}}$. With the assumption that the external nutrient concentrations, $c_{ext}$, are saturating with respect to the $K_M$ of the nutrient transporters, this simplifies to $\nu_{\max}$ giving us:

$$\frac{dc_{pc}}{dt} = \overbrace{\nu_{\max}\phi_M}^{\text{precursor synthesis rate}} - \gamma(c_{pc})\phi_R^{\text{free}} - c_{pc}\lambda$$
$$\underbrace{\phi_M}_{\text{Metabolic mass fraction}} = 1 - \phi_R(t) - \underbrace{\phi_0}_{\text{Constitutive protein mass fraction}} - \phi_{AR}(t) \tag{S6}$$

Conceptually, one could divide the proteome into as many subsectors as there are proteins, but we choose to focus on this limited set to simplify our model. The ribosomal and antibiotic-resistance fractions are easy to identify from experimental data and are clearly linked to a physiological role. For the purpose of our model, the remaining proteins can be divided into two fractions, a constitutive fraction that is not differentially regulated across the conditions we consider, and a metabolic fraction that has a characterizable rate, $\nu$, with which the metabolic proteins generate precursors.

If we solely consider steady-state growth rates, there is a degeneracy between $\phi_0$ and $\nu$ in our model: higher $\phi_0$ and consequently lower $\phi_M$ can be tuned to achieve the same growth rate by increasing $\nu_{\max}$. We parametrize $\phi_0$ by observing the amount of free proteome available in over-expression experiments [20, 21], which in turn allows us to define $\nu$ for a given media from observed steady-state growth. While this operationalizes our parameter, the so-called metabolic proteins likely include proteins that do not participate in metabolism. It might be more accurate to consider the metabolic proteins as "counter-ribosomal" proteins that are regulated in opposition to ribosomal proteins in the stringent response. The degeneracy between $\phi_0$ and $\nu$ can also be broken by observing the dynamic response to changing conditions.

By explicitly modeling the dynamics of antibiotics entering the cells and inhibiting ribosomes we can compare the relative speeds of environmental change and protein synthesis and quantify the relative fitness of different acclimatory strategies. The intracellular antibiotic concentration is primarily determined by the balance between influx from the environment, the degradation by antibiotic resistance proteins and dilution due to cellular growth [5, 12, 22–25]. The first two processes are characterized by $k_p$, the permeation rate through the membrane, and $k_{cat}$, the Michaelis-Menten binding of antibiotic-resistance proteins. The binding and unbinding of antibiotics from ribosomes also affects the intracellular concentration, especially during dynamic shifts. The intracellular concentration of antibiotics (chloramphenicol), $C_{in}$, is:

$$\frac{d\,C_{in}}{dt} = k_p \cdot \overbrace{\Big( C_{out}(t) - C_{in}(t) \Big)}^{\text{Environmental Coupling}} - \overbrace{\frac{k_{cat} n_{AR} \phi_{AR} C_{in}}{C_{in} + K_m^{AR}}}^{\text{Deactivation by Proteins}} + \underbrace{k_{off} \cdot C_{bnd} - k_{on} \cdot C_{in} \cdot (\phi_R n_R - C_{bnd})}_{\text{(Un)Binding from Inhibited Proteins}} - \lambda C_{in} \tag{S7}$$

where $C_{out}$ is the external concentration of antibiotics, $C_{bnd}$ is the concentration of antibiotics bound to ribosomes, and the rest of the numeric parameters are given with their citations in Table 1. The concentration of unbound ribosomes is given by $C_R^{\text{free}} = \phi_R n_R - C_{bnd}$, where the factor $n_R$ converts from mass fraction of proteins to the volumetric concentration in the cell:

$$C_X = \frac{\rho_{\text{cell}} \cdot f_{\text{protein}}}{m_{\text{X}} N_A} \cdot \phi_X = n_X\, \phi_X \tag{S8}$$

In the above, $m_X$ is the molar mass of the given protein. In equations S1 and S6 above,

| Symbol | Description | Value | Units | Reference |
|---|---|---|---|---|
| $\gamma_{\max}$ | max translation rate | 9.653 | 1/hr | Scott et al., 2010 |
| $\nu_{\max}$ | max precursor generation rate | 5 | 1/hr | This study, Dai et al. 2016, Scott et al. 2014 |
| $K_M$ | Binding constant (precursors to ribosome) | 0.03 | - | Chure & Cremer 2023 |
| $\phi_0$ | Allocation to other proteins | 0.65 | - | Dong et al 1995, Scott et al. 2014 |
| $k_p$ | Permeability | 800 | 1/hr | This study, Wu et al 1996 |
| $k_{\text{cat}}$ | CAT catalytic rate | 1.26e5 | 1/hr | Kleanthous and Shaw 1984 |
| $k_{\text{on}}$ | Binding rate | 2.31e6 | $1/(\text{hr} \cdot \text{M})$ | Harvey and Koch 1980, Hu et al 2020 |
| $k_{\text{off}}$ | Unbinding rate | 3.78 | $1/(\text{hr} \cdot \text{M})$ | Harvey and Koch 1980, Hu et al 2020 |
| $n_R$ | Ribosomal mass fraction-to-concentration factor | 63.7 | $\mu$M | This study |
| $n_{\text{CAT}}$ | CAT mass fraction-to-concentration factor | 536 | $\mu$M | This study |
| $K_M^{\text{ABX}}$ | Binding constant (antibiotics to ribosome) | 15e-6 | $\mu$M | Hu et al 2020 |

TABLE S1. Model parameters

$\phi_R^{\text{free}} = \frac{C_R^{\text{free}}}{n_R}$. Completing the picture, the bound antibiotic concentration is given by:

$$\frac{d\, C_{bnd}}{dt} = k_{on} \cdot C_{in} \cdot (\phi_R n_R - C_{bnd}) - k_{off} \cdot C_{bnd} - \lambda C_{bnd} \tag{S9}$$

The differential equations were coded in Julia [26] using the DifferentialEquations [27, 28] package to solve the equations and the JuMP[29] package for certain optimizations using the equations.

##### 1. *Model Parameters*

### B. (Outer) Membrane Permeability of Gram-negative Bacteria

For antibiotics to act they must first be transported into a cell by crossing the cell membrane. In the case of Gram-negative bacteria, this typically means crossing the outer and inner membrane. The literature has focused primarily on the permeability of the outer membrane (possibly because because the selective permeability of the Gram-negative outer membrane has posed an ongoing challenge to drug development and because once antibiotics can typically cross the inner membrane if they weren't excluded by the outer membrane.) When the outer membrane is damaged/stripped away with EDTA, Gram-negative bacteria are more easily druggable, and this was taken as evidence that the outer membrane is the primary barrier [30]. Molecules that are smaller than 600 Da and hydrophilic can diffuse into *E. coli* via OmpF [31, 32]. Similar exclusion criteria are true in other Gram-negative bacteria [33, 34]. More detailed "eNTRy" rules have since been developed as a heuristic for describing in greater detail which characteristics are associated with improved intracellular accumulation [35]. The literature primarily focuses on structural and molecular mechanisms of permeation and resistance, rather than the kinetics of the processes.

Nonetheless, there are some quantifications of permeation. The range of different antibiotic permeabilities is dizzyingly large – it spans from $10^{-8}$ to $10^{2}$ cm/s [36]. However, we can do a lot better by motivating our model based on small molecule antibiotics like chloramphenicol. Abdel Sayed [37] measured the accumulation of chloramphenicol in *E. coli* but the dosages are preposterously high (starting at $\sim$ 30x MIC and going up [38]) so they are not relevant to our physiological study. Mortimer and Piddok [39] measured the accumulation rates of chloramphenicol in *E. coli* from which the permeation rate might be inferred. However, the extensive body of work describing the relationship between molecular characteristics and their outer membrane permeability allows us to infer the permeation rate of chloramphenicol based on its hydrophobicity [31]. Based on a log10(octanol) partition coefficient of 1.02-1.07 [40, 41], chloramphenicol is expected to have a permeability of $\sim 10^{-5}$ cm/s. Converting to the whole-cell permeability rate (measured in units of 1/s) requires multiplying by the surface area to volume ratio of *E. coli* which ranges from 4-8 $\mu$m$^{-1}$ [42, 43]. The whole-cell permeability is then $\sim 0.6s^{-1}$.

#### C. Intracellular Antibiotics Dynamics

As discussed in Section S2 B, Gram-negative bacteria have a first line of defense to antibiotics through the limited permeability of their outer membrane. Once antibiotics have entered the cell efflux pumps and various antibiotic-inactivating enzymes work to minimize the effect of antibiotics on cellular funcction. There are two important types of important antibiotic reactions in the cell: the binding and unbinding of the antibiotic to its target and the resistance mechanism that impedes the antibiotic.

To model the resistance mechanism, we consider the effects of an enzyme that inactivates antibiotics, motivated by the example of chloramphenicol acetyltransferase. Our model does not consider the cost of consuming 1-2 acetyl groups to deactivate the chloramphenical, though this might be significant in resource-limited scenarios. We expect our model to also work for describing efflux pumps when the energetic cost[44] of running the pump is comparatively small and when the extracellular concentration is not significantly affected by the activity of the pump. In wild-type *E. coli* MG-1655 with chloramphenicol acetyltransferase (CAT), knocking out the AcrAB efflux pump noticeably affect resistance [45]. We surmised that in the presence of CAT and efflux pumps, CAT is primarily responsible for chloramphenicol resistance.

#### 1. *Catalytic Rate of Chloramphenicol Acetyltransferase*

The chloramphenicol acetyltransferase (CAT) catalyzed reaction consumes both chloramphenicol and acetyl-CoA. The complete reaction mechanism includes a reaction pathway to a diacetylation of chloramphenicol, but it occurs much more slowly than the primary acetylation and is not necessary for understanding the essential physiological function of CAT [46]. The essential reaction can be modelled as a Michaelis-Menten reaction [47]. Though the reaction also depends on concentration of acetyl-CoA, for simplicity we consider the concentration of acetyl-CoA to be fixed. More precisely, acetyl-CoA varies from $\sim 150 - 450\mu$M [48]. Over the same range, the $k_{cat}$ of CAT varies between $30 - 45s^{-1}$ [47].

#### 2. *Binding and Unbinding of Chloramphenicol to Ribosomes*

The binding and unbinding of chloramphenicol from ribosomes has been previously characterized both in vivo and in vitro in assays with *E. coli* A19 and ML30 [49]. These parameters were previously used to constrain another model of cell growth focusing on protein synthesis [50]. Notably, the authors of the original binding data note that the agreement between in-vitro and in-vivo data "suggests that diffusion of [chloramphenicol] across the cell membrane is not a significant factor in the development of inhibition and that the intracellular concentration of free [chloramphenicol] rapidly becomes equal to the extracellular concentration." This hypothesis is further supported by our modelling of chloramphenicol diffusion into *E. coli*.

#### *3. Parameters for Other Antibiotics*

For concreteness, we chose to focus on chloramphenicol inhibition in *E. coli*. However, more broadly antibiotic permeation and binding rates vary across many orders of magnitude and as a result. The $K_D$ for quinolones varies from about $10^{-4}$ to $10^{-8}$ M [51]. Furthermore, the binding rates for a quinolone resistant Salmonella arising from a point mutant was 100x less than the wild-type [52]. The beta-lactams typically bind to penicillin-binding proteins irreversibly, so there is no $k_{off}$.

In spite of the broad distribution of antibiotic mechanisms and rates, two patterns are widely true. First, bacteria negotiate a tradeoff between more resistance and faster growth. Second, protein synthesis lags behind the permeation and binding of antibiotics. As a consequence, although exact antibiotic rates vary widely, we expect the phenomenon of anticipatory pre-synthesis of proteins to be widely relevant.

#### *4. Clinical and Environmental Concentrations of Antibiotics*

For contextualizing the relevance of the antibiotic concentrations investigated, we briefly review the antibiotic concentrations bacteria might experience outside of the lab. The recommended working concentration for chloramphenicol when preparing lab cultures is 25 $\mu$g/mL ($\sim 80\mu$M), which is about 5x the MIC.

Due to the prevalence of antibiotic use[53–55], microbes likely experience antibiotic exposures for extended durations. In agricultural soils, especially around concentrated animal feeding operations, there are persistent levels of residual antibiotics ($\sim$pM) in the soil [55]. Microbes in sewage treatment plants also likely experience persistent but low concentrations of antibiotics.

In hospitals, the concentrations are much higher. The highest doses of antibiotics experienced by bacteria are likely during sepsis and adjacent to topically applied antibiotic ointments. In clinical settings, antibiotics have to be administered at especially high concentrations to ensure the drug reaches the infected site at high enough concentrations. In sepsis, antibiotics are administered at extremely high doses periodically, as high as 100x the

MIC [56–59] to ensure the concentration stays high enough for the duration of the dosing period [60]. As a result, septic bacteria experience fluctuating antibiotics with extremely high peaks. Topical ointments can have even higher absolute concentrations because their toxicity is constrained by their limited diffusion. Neosporin, a common topical ointment, has a concentration of 11 mM cumulatively of bacitracin, neomycin and polymyxin which is >1000x MIC.

---


*

†



s41467-025-57789-4. Publisher: Nature Publishing Group.

10.1038/nmicrobiol.2016.231.

nlm.nih.gov/pmc/articles/PMC4380930/.

9193. doi:10.1128/jb.153.1.241-252.1983.

[40] Chemspider. (-)-chloramphenicol. URL https://www.chemspider.com/Chemical-Structure.5744.html?rid=f9224127-d167-47d1-a259-3d5acec9ef9d.

[41] Changhao Wu, Paul Clift, Christopher H. Fry, and John A. Henry. Membrane action of

chloramphenicol measured by protozoan motility inhibition. *Arch. Toxicol.*, 70(12):850–853, October 1996. ISSN 1432-0738. doi:10.1007/s002040050349.

[42] F. J. Trueba and C. L. Woldringh. Changes in cell diameter during the division cycle of Escherichia coli. *J. Bacteriol.*, 142(3):869, June 1980. doi:10.1128/jb.142.3.869-878.1980.

[43] O. Pierucci. Dimensions of Escherichia coli at various growth rates: model for envelope growth. *J. Bacteriol.*, 135(2):559–574, August 1978. ISSN 0021-9193. doi:10.1128/jb.135.2.559-574.1978.

[44] Note1. The efflux pumps operate by using the proton gradient across the inner membrane, at the opportunity cost of using that same gradient for turning an ATPase.

1553-7358. doi:10.1371/journal.pcbi.1008106.

[52] Hassan M. Al-Emran, Anke Heisig, Denise Dekker, Yaw Adu-Sarkodie, Ligia Maria Cruz Espinoza, Ursula Panzner, Vera von Kalckreuth, Florian Marks, Se Eun Park, Nimako Sarpong, Jürgen May, and Peter Heisig. Detection of a Novel gyrB Mutation Associated With Fluoroquinolone-Nonsusceptible Salmonella enterica serovar Typhimurium Isolated From a Bloodstream Infection in Ghana. *Clin. Infect. Dis.*, 62(Suppl):147–149, March 2016. ISSN 1537-6591. doi:10.1093/cid/civ790.

[53] Whocc. Defined Daily Dose - Definition and general considerations, February 2018. URL https://atcddd.fhi.no/ddd/definition_and_general_considera. [Online; accessed 29. Jul. 2024].

[54] Prevention WHO Surveillance and Control. Who report on surveillance of antibiotic consumption, 2019.

[60] Michiel Haeseker, Thomas Havenith, Leo Stolk, Cees Neef, Cathrien Bruggeman, and Annelies

Verbon. Is the standard dose of amoxicillin-clavulanic acid sufficient? *BMC Pharmacol. Toxicol.*, 15(1):1–8, December 2014. ISSN 2050-6511. doi:10.1186/2050-6511-15-38.
